# FicD Sensitizes Cellular Response to Glucose Fluctuations in Mouse Embryonic Fibroblasts

**DOI:** 10.1101/2024.01.22.576705

**Authors:** Burak Gulen, Lisa N. Kinch, Kelly A. Servage, Aubrie Blevins, Nathan M. Stewart, Hillery F. Gray, Amanda K. Casey, Kim Orth

## Abstract

During homeostasis, the endoplasmic reticulum (ER) maintains productive transmembrane and secretory protein folding that is vital for proper cellular function. The ER-resident HSP70 chaperone, BiP, plays a pivotal role in sensing ER stress to activate the unfolded protein response (UPR). BiP function is regulated by the bifunctional enzyme FicD that mediates AMPylation and deAMPylation of BiP in response to changes in ER stress. AMPylated BiP acts as a molecular rheostat to regulate UPR signaling, yet little is known about the molecular consequences of FicD loss. In this study, we investigate the role of FicD in mouse embryonic fibroblast (MEF) response to pharmacologically and metabolically induced ER stress. We find differential BiP AMPylation signatures when comparing robust chemical ER stress inducers to physiological glucose starvation stress and recovery. Wildtype MEFs respond to pharmacological ER stress by downregulating BiP AMPylation. Conversely, BiP AMPylation in wildtype MEFs increases upon metabolic stress induced by glucose starvation. Deletion of FicD results in widespread gene expression changes under baseline growth conditions. In addition, FicD null MEFs exhibit dampened UPR signaling, altered cell stress recovery response, and unconstrained protein secretion. Taken together, our findings indicate that FicD is important for tampering UPR signaling, stress recovery, and the maintenance of secretory protein homeostasis.

**Significance Statement:** The chaperone BiP plays a key quality control role in the endoplasmic reticulum, the cellular location for the production, folding, and transport of secreted proteins. The enzyme FicD regulates BiP’s activity through AMPylation and deAMPylation. Our study unveils the importance of FicD in regulating BiP and the unfolded protein response (UPR) during stress. We identify distinct BiP AMPylation signatures for different stressors, highlighting FicD’s nuanced control. Deletion of FicD causes widespread gene expression changes, disrupts UPR signaling, alters stress recovery, and perturbs protein secretion in cells. These observations underscore the pivotal contribution of FicD for preserving secretory protein homeostasis. Our findings deepen the understanding of FicD’s role in maintaining cellular resilience and open avenues for therapeutic strategies targeting UPR-associated diseases.

## Introduction

Cellular stress responses alter the balance of protein synthesis, modification, and degradation in the cell to maintain protein homeostasis and cellular function. The unfolded protein response (UPR) is an adaptive signaling pathway that helps to restore cellular homeostasis to varying levels of ER stress (*1*), ranging from mild to maladaptive (*2*). Activation of the UPR is regulated by the essential ER chaperone protein BiP (a.k.a. glucose regulated protein 78 or heat shock protein family A member 5 (Hspa5)) and results in the activation of a complex signaling network promoting both cell survival and apoptosis pathways (*3–5*). As the main chaperone residing in the ER, BiP plays a critical role in promoting both the correct folding and transport of newly synthesized proteins passing through the ER and the degradation of misfolded proteins by the ER associated protein degradation (ERAD) pathway (*1*). The UPR is activated when the ER protein folding capacity is insufficient to cope with the burden of unfolded proteins accumulating in the ER (*6*). UPR aims to restore homeostasis by attenuating global translation and transcription, and by enhancing the folding capacity of the cell through selective transcription and translation of chaperones like BiP (*7, 8*).

UPR is activated by the three transmembrane ER stress sensors that serve as distinct yet intertwined signaling branches: 1) PERK (protein kinase RNA (PKR)-like ER kinase), 2) IRE1α (inositol-requiring enzyme 1α), and 3) ATF6 (activating transcription factor 6) (*1, 3*). When unfolded proteins accumulate in the ER lumen, BiP dissociates from the three signal transducers to implement the UPR (*1*) (*4*) (**Fig. 1A**). PERK phosphorylates eIF2α to repress protein synthesis and induce preferential translation of Atf4, the master transcription factor of the integrated stress response. Ire1α processes unspliced *X-box binding protein 1* (*XBP1*) mRNA, promoting the translation of the XBP1 transcription factor, which upregulates ER chaperones and ERAD components. Membrane bound Atf6 translocates from the ER to the Golgi apparatus, where it is processed into a cytoplasmic transcription factor that upregulates ER chaperones and lipid synthesis. The regulation of transcription during UPR is critical for normal cellular function and health; however, failure to restore homeostasis ultimately leads to a maladaptive/pathologic phase encompassing activation of pro-apoptotic genes and programmed cell death (**Fig. 1A**). Chronic or dysregulated UPR is associated with a variety of diseases (*9–11*).

**Fig. 1:**
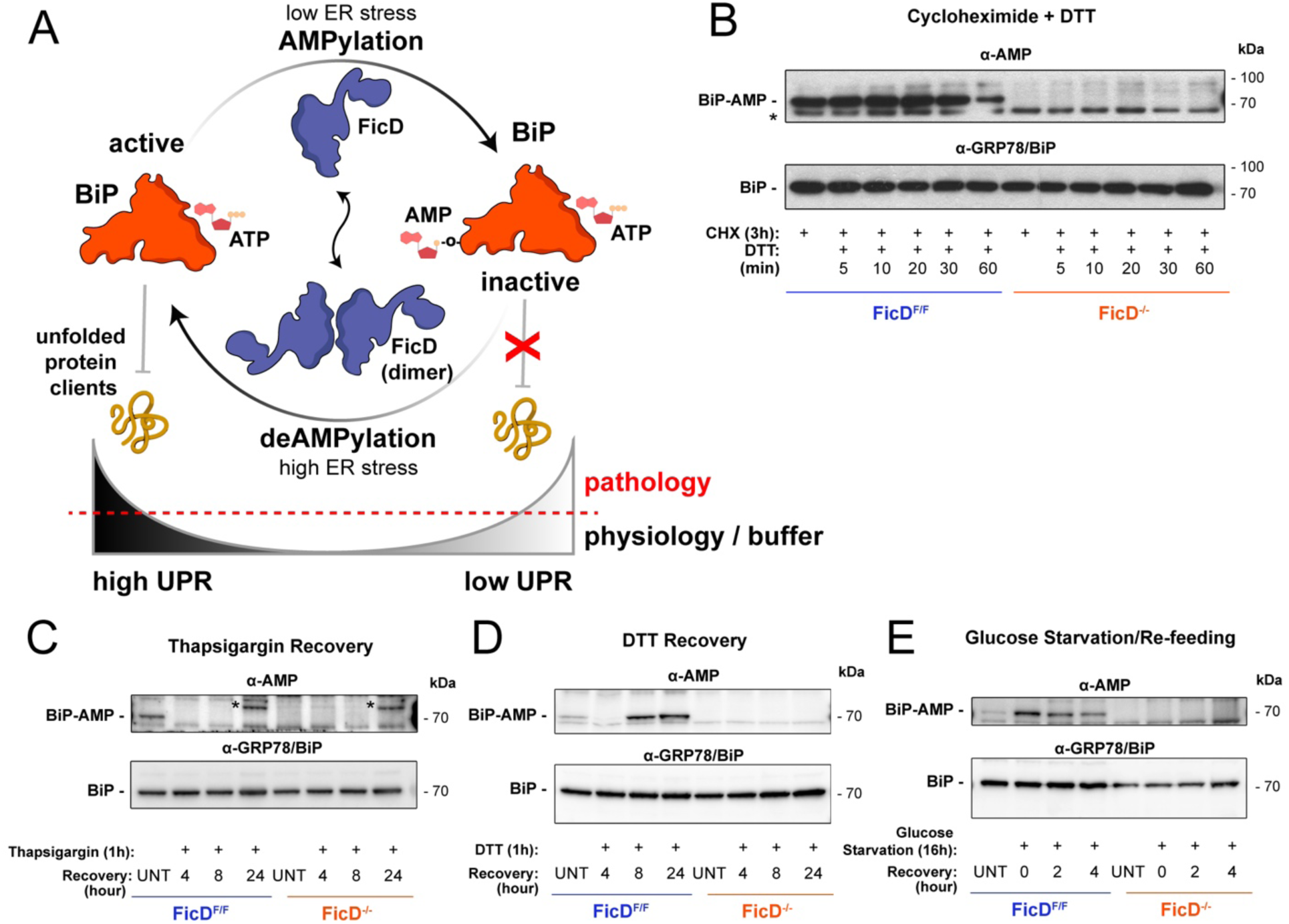
AMPylation of BiP and response to ER stress in mouse embryonic fibroblasts (MEFs). **A)** Regulation of BiP activity by AMPylation. Oligomeric state of FicD mediates the antagonistic activities of FicD (*62, 63*). **B)** AMPylation responds to ER stress in *FicD*^F/F^ MEFs and *FicD*^-/-^ MEFs. Western blots showing AMPylated and total BiP. After 3h of cycloheximide (CHX, 100 µg/ml) treatment. Adding DTT (1mM) in CHX treated MEFs increases the ER stress and thus decreases the AMPylation of BiP over time. *FicD*^-/-^ cells do not show BiP AMPylation in the absence of FicD. *unidentified band, present in both genotypes. **C-E)** Western blots showing AMPylated and total BiP. *unidentified band, present in both genotypes. MEFs are treated with ER stressing drugs **C)** thapsigargin (TG, 1µM) or **D)** DTT (1mM) for 1h or **E)** starved for glucose for 16h following the recovery by either removing the drug or adding glucose. UNT: untreated/unstressed.

AMPylation (i.e., adenosine monophosphate (AMP) transfer to proteins) is a post translational modification (PTM) conserved across all kingdoms of life. Catalyzed primarily by the large family of filamentation induced by cyclic-AMP (Fic) enzymes (*12–19*), AMPylation modulates cellular functions (*20*), typically resulting in inactivation of target proteins. Metazoans carry a single Fic domain protein encoding gene (*FicD*). *FicD* encodes a bifunctional enzyme localized in the endoplasmic reticulum (ER) membrane with a luminal catalytic domain (*21*) that reversibly AMPylates or deAMPylates the ER chaperone BiP (*22, 23*). In a state of homeostasis, FicD acts as an AMPylator, generating an inactive reserve pool of the BiP chaperone within the ER lumen (*24–26*). During ER stress, de-AMPylation of BiP by FicD re-activates this pool of chaperones to aid in resolving stress induced by unfolded proteins (*27*).

Alternate proposals exist in the field pertaining to the fitness benefits of FicD-dependent BiP AMPylation. Some studies anticipate that AMPylation of BiP acts as a molecular rheostat for UPR. The rheostat allows different cells to respond to varying ER stress thresholds by maintaining excess BiP in a reversibly inactive state (*12, 13, 22, 27*). Previous work using animal models support this hypothesis, as loss of FicD results in aberrant UPR signaling in tissues and sensitizes tissues to damage in *Drosophila*, *C. elegans*, and mice (*28–31*). In the absence of FicD, tissues facing repetitive stress display increased damage and delayed recovery after each insult. AMPylation and rapid inactivation of BiP have also been suggested to benefit cells by preventing over chaperoning and excessive ERAD pathway activation (*32*). This hypothesis is supported by AMPylation profiles of BiP in cells that correlate with UPR activation and by in vitro kinetic modeling of protein-folding homeostasis in the ER with and without a reserve source of BiP for fast reactivation (*22, 26, 33*). Despite these previous studies, key questions remain regarding how specific cellular stresses induce changes in BIP activity.

We sought to better define the fitness role of FicD in mammalian cells experiencing physiological and pharmacological-induced ER stress. We recently produced a floxed, Flag-tagged *FicD* allele in mice to study the effect of FicD in response to ER stress and found that loss of FicD leads to elevated UPR and reduced recovery from ER stress (*30*). Here, we isolated and immortalized control Flag-tagged FicD mouse embryonic fibroblasts (MEFs) and FicD knockout MEFs (*FicD*^F/F^ MEFs and *FicD*^-/-^ MEFs, respectively) (**Fig. S1**) to characterize the role of FicD in response to ER stress at a cellular level. Using a variety of methods, we report that the absence of FicD causes fundamental changes in the transcriptome, leading to increased expression and secretion of extracellular matrix (ECM) proteins. The *FicD*^-/-^ MEFs lack a transcriptional response to glucose starvation and display a dampened transcriptional UPR. Taken together, our data support the hypothesis that AMPylation of BiP tempers the activation of the UPR response under physiological ER stress inducing conditions.

## Results

### MEFs undergo reversible AMPylation of the ER chaperon BiP

To study the cellular response to BiP AMPylation, FLAG-tagged *FicD*^F/F^ MEFs and *FicD*^-/-^ mutant MEFs were isolated, cultured and immortalized by transfection with SV40 antigen containing plasmid. While the limited expression levels of FicD did not allow its endogenous detection through western blots, FicD can be monitored both at the transcript and enzymatic activity level. Using both RT-qPCR of *FicD* transcript and assessment of BiP-AMPylation levels via Western blot analysis, we observed that immortalized *FicD*^F/F^ MEFs grown in standard growth media produce *FicD* mRNA and functional FicD enzyme capable of reversibly AMPylating BiP, whereas the *FicD*^-/-^ MEFs do not express *FicD* or AMPylate BiP (**Fig. 1B** and **Fig. S2**).

Previously, we have shown that CHX treatment, which inhibits mRNA translation, enhances BiP AMPylation in cell lines (*22, 29*). To determine how loss of FicD influences this response, we treated our immortalized *FicD*^F/F^ and *FicD*^-/-^ MEFs with CHX. Consistent with previous observations, BiP AMPylation increases in *FicD*^F/F^ MEFs following CHX exposure as monitored by western blots using monoclonal α-AMP antibody. Unlike control cells, *FicD*^-/-^ MEFs exhibited no detectable BiP AMPylation (**Fig. 1B**) (*22*). Next, we perturbed the ER’s oxidation state in CHX-treated cells by introducing the reducing agent DTT into the culture media. This treatment results in the reversal of BiP AMPylation in *FicD*^F/F^ MEFs over time (60 minutes; **Fig. 1B**), indicating that BiP AMPylation levels change in response to different cellular stresses, even when global protein synthesis levels are inhibited.

Next, we surveyed the AMPylation status of BiP during recovery from ER stress caused by treatment with thapsigargin (TG) or DTT alone. TG inhibits the sarco/ER Ca²⁺ ATPase, thereby perturbing Ca^2+^ signaling in the ER (*34*). Following exposure to either TG or DTT, MEFs were allowed to recover from ER stress in fresh media. Unstressed cells exhibit a baseline AMPylation level of BiP, which disappears in response to both pharmacological ER stress treatments (**Fig. 1C and D**). Irreversible TG-mediated stress results in a loss of BiP AMPylation that persisted even after 24-hours of recovery. By contrast, MEFs recovering from reversible DTT-mediated ER stress display a reemergence of BiP AMPylation at around 8 hours (**Fig. 1D**). The protein levels of BiP do not change significantly under these conditions (**Fig. 1C and D**).

In addition to these pharmacological stressors, we sought to study a physiological stress condition in MEFs by depriving cells of glucose in the growth media. Glucose starvation is an established physiological stress that leads to induction of the UPR (*2, 35, 36*). Interestingly, glucose starvation in *FicD*^F/F^ MEFs resulted in a behavior like that observed upon CHX treatment, boosting BiP AMPylation. After adding back glucose to the media, the AMPylation levels decreased towards the baseline exhibited by unstressed cells (**Fig. 1E**).

### *FicD***^-/-^** MEFs exhibit altered UPR gene expression patterns during stress and recovery

To complement our analysis of BiP AMPylation during an ER stress, we used RT-qPCR to measure how mRNA levels of UPR genes change during various stress-inducing conditions in the *FicD*^F/F^ and *FicD*^-/-^ MEFs. We predicted that UPR signaling and recovery in *FicD*^F/F^ and *FicD*^-/-^ MEFs may be differentially altered under these various conditions in the presence and absence of BiP AMPylation (**Fig. 1C-E**)

Treatment with TG for over one hour increased the levels of *Atf3* and s*Xbp1* transcripts in *FicD*^F/F^ MEFs, and their relative expression was significantly higher in *FicD*^-/-^ MEFs across all time points. In contrast, relative expression levels of *Chop/Ddit3* and *Atf4* increased to the same extent in both genotypes (**Fig. 2A** and **Fig. S2A**). Additionally, expression patterns of the *BiP/Hspa5* transcript differed significantly between *FicD*^F/F^ and *FicD*^-/-^ MEFs during TG treatment. Over the one hour treatment with TG, expression levels of the *BiP/Hspa5* transcript steadily increased in *FicD*^F/F^ MEFs. However, in *FicD*^-/-^ MEFs expression levels of the *Bip/HspA5* transcript were significantly diminished within the first 15 minutes of treatment and remained significantly diminished during the one hour TG treatment (**Fig. S2A**). This data supports the proposal that induction of UPR by acute TG treatment was differentially regulated in *FicD*^F/F^ and *FicD*^-/-^ MEFs.

**Fig. 2:**
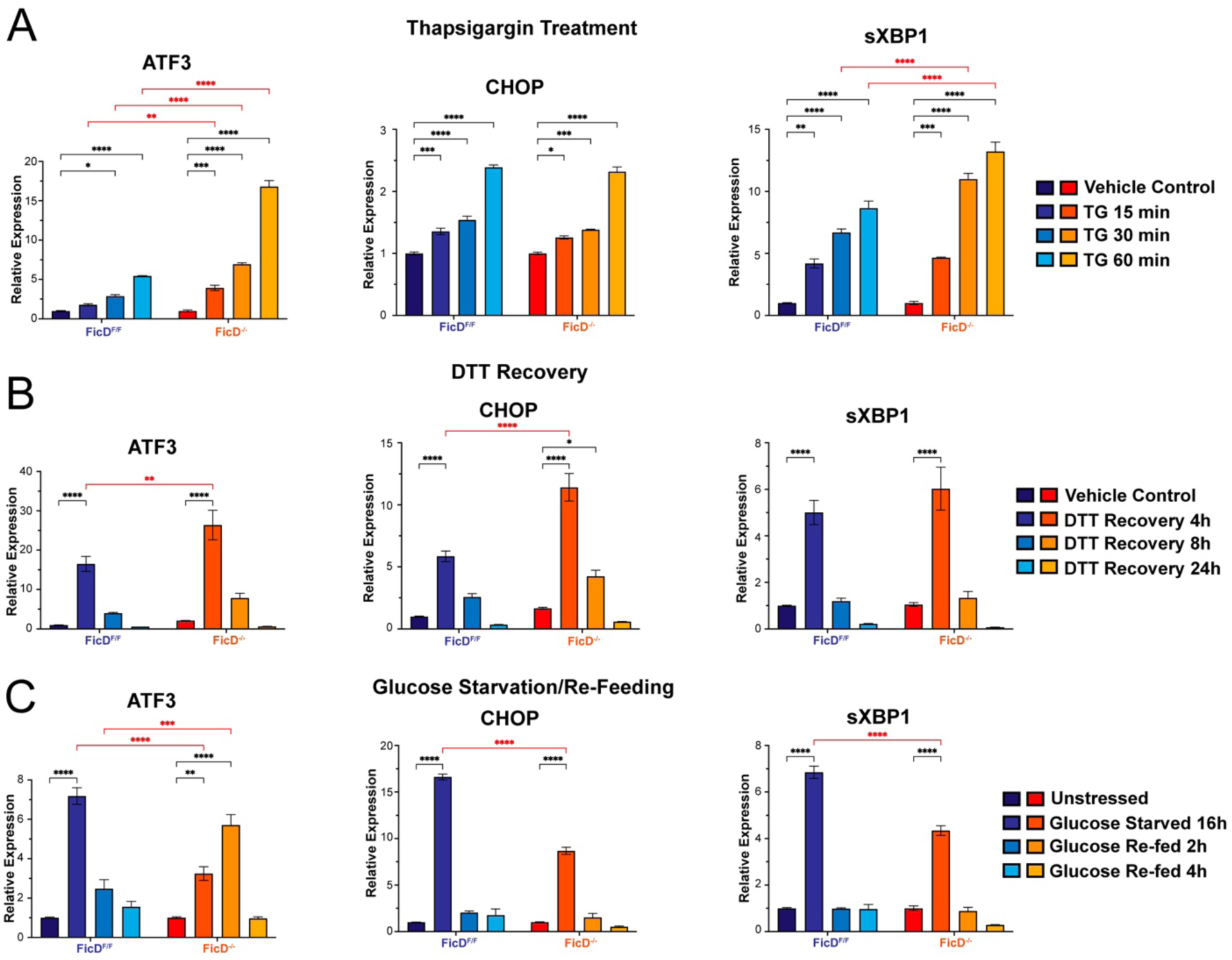
Thapsigargin (TG), DTT, and Glucose Starvation induced ER stress of *FicD*^F/F^ and *FicD*^-/-^ MEFs. RT-qPCR showing relative expression levels of UPR marker genes ATF3, CHOP, and spliced XBP1 (sXBP1) upon **A)** treatment of MEFs with Thapsigargin (1µM), or **B)** recovery of MEFs from 1h DTT (5mM) exposure, or **C)** glucose starvation (for 16h) and re-feeding of MEFs for indicated time points. Error bars represent the standard deviation of 3 biologically independent repeat of the experiment with 4 technical replicates. Genotypes are demonstrated by shades of blue for *FicD*^F/F^ and shades of orange for *FicD*^-/-^ MEFs. Two-way ANOVA with Tukey multiple comparison test is applied to determine the significance. P values: 0.1234 (ns), 0.0332 (*), 0.0021 (**), <0.0001 (****). Non-significant comparisons are not shown for clarity.

We next analyzed how *FicD*^F/F^ and *FicD*^-/-^ MEFs recovered from stress by treating cells with DTT followed by washing and observing gene expression after 4, 8 and 24-hours of recovery. The mRNA for *Atf3*, *Chop/Ddit3* and s*Xbp1* decreased over time to baseline levels (after 24-hours of recovery). In the early (4-hour) recovery timepoint, the levels of *Atf3*, *Chop/Ddit3*, and *BiP/HspA5* transcripts were significantly higher in *FicD*^-/-^ cells, suggesting a delayed recovery from UPR in *FicD*^-/-^ MEFs (**Fig.2B, Fi. S2B**). Changes in *Atf4* transcript responded similarly in both genotypes (**Fig. S2B**). Thus, with the exception of the *Atf4* transcript, the relative expression levels of UPR genes increased in response to pharmacological induction of ER stress in the *FicD*^-/-^ MEFs.

Finally, we analyzed how *FicD*^F/F^ and *FicD*^-/-^ MEFs responded to and recovered from a physiological stress, glucose starvation. The relative expression levels of *Atf3*, *Chop/Ddit3*, s*XBP1, Atf4,* and *FicD* were significantly increased in *FicD*^F/F^ MEFs upon starvation and returned to near basal levels after refeeding (**Fig. 2C, Fig. S2C**). The relative expression levels of *BiP/HspA5* were also significantly increased in the *FicD*^F/F^ MEFs but did not return to basal levels within 4-hours of recovery (**Fig. S2C**). The response to this metabolic stress in *FicD*^-/-^ MEFs was dampened, with none of the transcripts elevated to the same extent during glucose starvation. Upon refeeding the *FicD*^-/-^ MEFs, the relative expression levels of *Chop/Ddit3*, s*Xbp1*, and *BiP/HspA5* decreased to basal levels, with patterns similar to those observed in *FicD*^F/F^ MEFs (**Fig. 2C, Fig. S2C**). Resembling the recovery from DTT-mediated ER stress, *Atf4* levels changed similarly in both genotypes during glucose starvation and refeeding (**Fig. S2C**). However, a distinct expression pattern was observed for *Atf3* transcripts in *FicD*^-/-^ MEFs. After 2-hours of refeeding, the *Atf3* transcript level was elevated before lowering back towards basal levels at 4-hours of refeeding (**Fig. 2C**). Taken together, relative UPR expression analysis of *FicD*^F/F^ and *FicD*^-/-^ MEFs indicates differential regulation under TG, DTT, and glucose starvation conditions, and loss of FicD had significant but distinctive effects under each condition.

### Loss of FicD induces dramatic changes in gene expression profiles of MEFs

We were intrigued by our findings in *FicD*^F/F^ MEFs that showed increased BiP AMPylation correlated with strong UPR activation during the physiological stress of glucose starvation (**Fig. 1E**, **Fig. 2C**). These observations directly conflict with previous reported AMPylation profiles of BiP and accepted models of FicD catalytic activity for deAMPylation of BiP during a pharmacologically induced stress (*22, 26–30, 33*). In addition, altered UPR gene transcript levels during starvation and recovery in *FicD*^-/-^ MEFs compared to *FicD*^F/F^ MEFs suggested that loss of FicD activity altered the response of these cells to this physiologically relevant stress.

To investigate whether additional molecular pathways could be altered in *FicD*^-/-^ MEFs when compared to *FicD*^F/F^ MEFs under glucose starvation, we performed RNAseq. Data was collected for *FicD*^F/F^ and *FicD*^-/-^ MEFs treated with four different conditions: unstressed (standard growth media), 18 hours of glucose starvation; 2-hours of glucose refeeding; and 4-hours of glucose refeeding (**Fig. 3A**). Principal component analysis (PCA) of all gene counts cluster the biological replicates from each condition, while the clusters for genotypes and treatments segregate. The *FicD*^F/F^ and *FicD*^-/-^ MEF transcriptomes exhibit similar trends relative to metabolic stress treatments. The gene expression response to glucose starvation changes the least compared to unstressed cells. Glucose refeeding at 2-hours initiates the largest transcriptome response, and 4-hours refeeding trends back towards the unstressed state. Notably, the largest spread of the gene expression data (indicated by PC1) results from a single change, the absence of FicD (**Fig. 3B**). We compared RNA-seq reads from various pairwise conditions to better understand the transcriptome changes associated with 1) metabolic stress treatment and recovery in *FicD*^F/F^ 2) metabolic stress treatment and recovery in *FicD*^-/-^ MEFs and 3) genotype differences in unstressed and starved MEFs (**Fig. 3C**).

**Figure 3.**
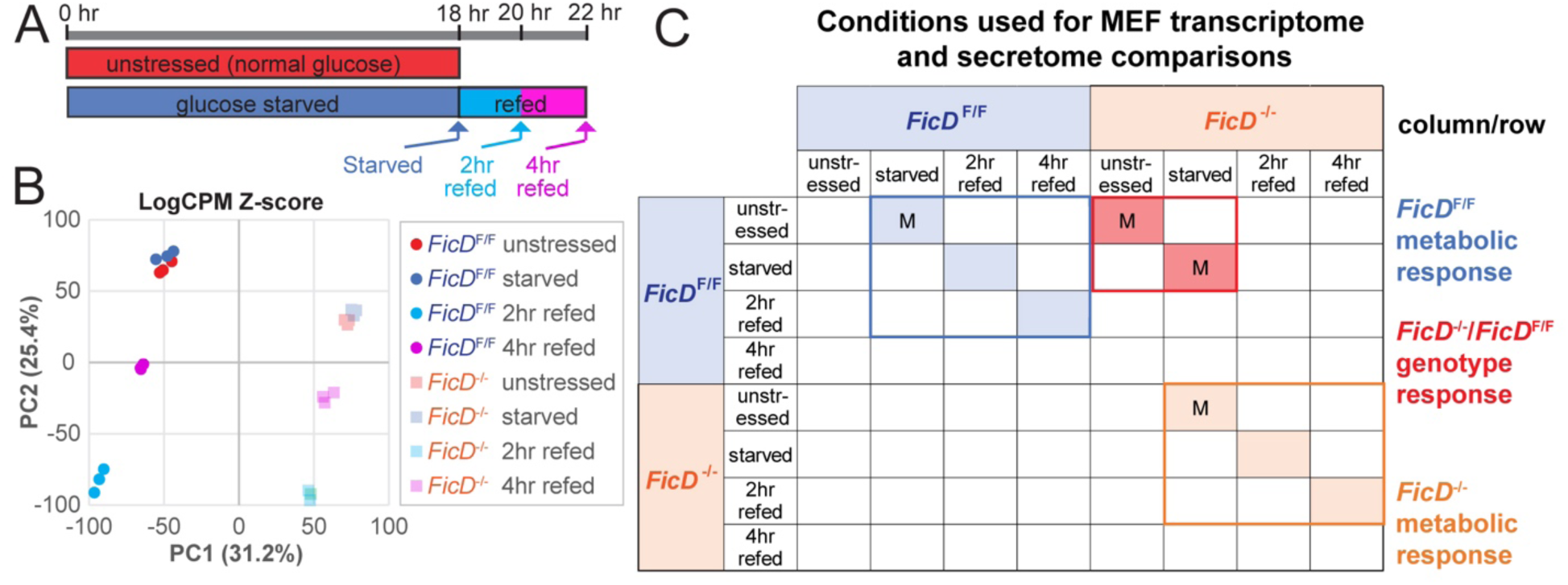
MEF genotype and metabolic treatment comparisons. **A)** Timeline of MEF treatments. **B)** PCA plot of Log2CPM Z-scores for indicated genotypes and treatments in triplicate. **C)** DEG were calculated by comparison of conditions indicated in rows with respect to conditions in columns. Comparisons indicated by “M” were also compared by MS/MS of secreted proteins.

### Glucose starvation induces PERK responsive UPR that recovers with refeeding

Differentially expressed genes (DEGs) were defined for *FicD*^F/F^ MEFs subjected to the various treatments (**Fig. 3A and C**, *FicD*^F/F^ metabolic response). The *FicD*^F/F^ glucose-starved cells exhibited 150 significantly upregulated genes compared to unstressed MEFs, including eight elevated genes classically related to UPR (**Fig. 4A**). The starvation responsive UPR genes from the *FicD*^F/F^ MEFs include those belonging to the PERK modulated cascade leading to apoptosis (*Atf4*, *Atf3*, *Chop/Ddit3*, *Chac1*, *Ero1a*), consistent with our RT-qPCR measurement observed in **Figure 2**.

**Figure 4.**
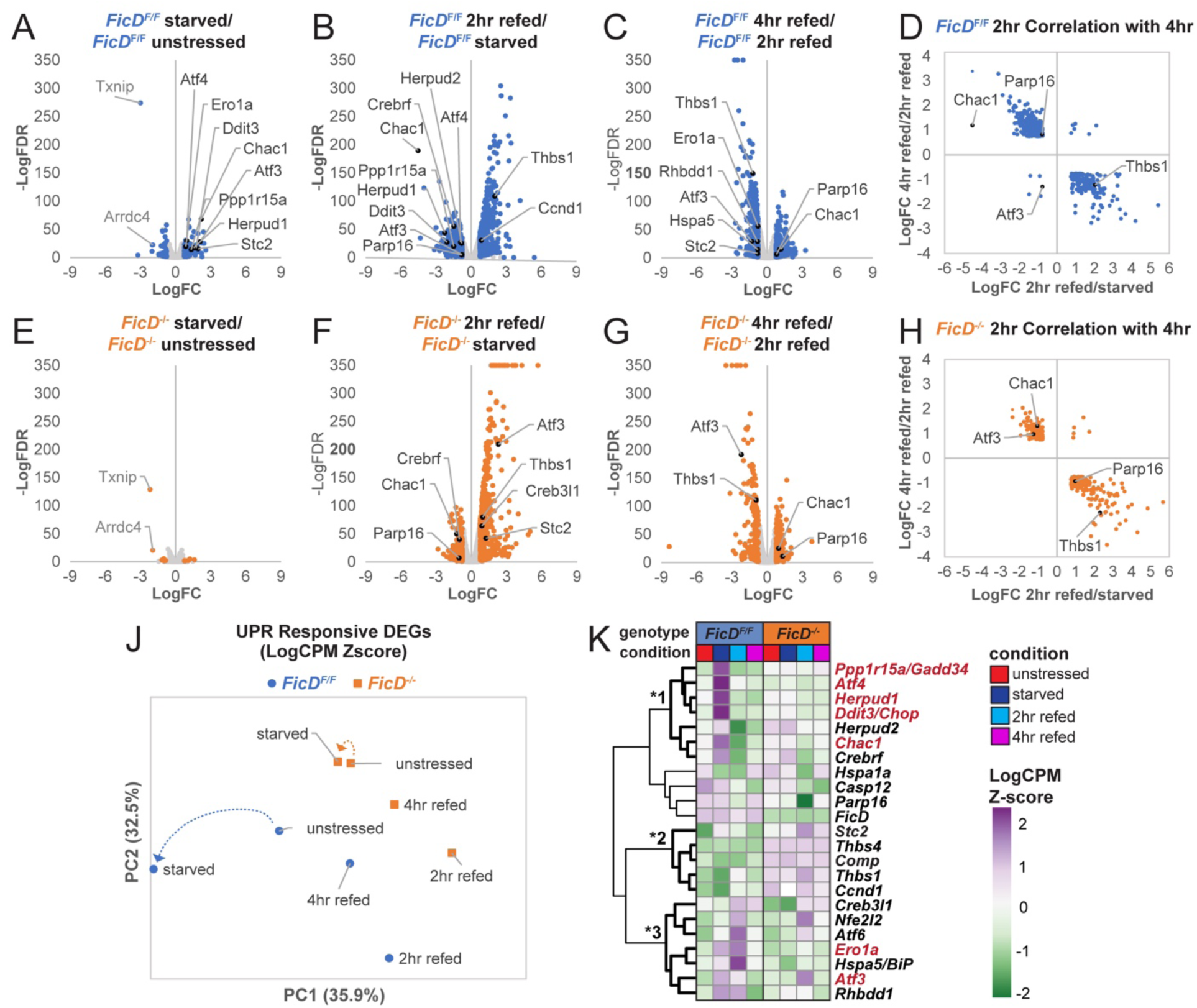
*FicD*^-/-^ MEFs exhibit muted response to homeostatic metabolic stress. **A-C)** *FicD*^F/F^ and **E-G)** *FicD*^-/-^ volcano plots highlight RNA-seq comparisons for conditions labeled above graphs, with nodes representing DEGs (blue) or other genes (gray). UPR DEGs (black) and common downregulated genes in starved cells (gray) are labeled. **D)** *FicD*^F/F^ and **H)** *FicD*^-/-^ LogFC inverse correlation between overlapping DEGs for 2-hour refed/starved and 4-hour refed/2-hour refed, with UPR genes labeled. **J)** PCA plot with arrows pointing towards starved conditions from unstressed conditions and **K)** heatmap of Log2CPM Z-scores for UPR DEGs, with three clusters indicated by * adjacent to bolded branches. PERK responsive UPR genes are labeled in red.

The role of the UPR in the *FicD*^F/F^ MEFs response to glucose starvation is further supported by enriched functions among the upregulated DEGs (**Table 1**, UPR terms marked by *). Both starvation responsive terms and UPR associated terms were enriched. In addition, functional overlap between glucose starvation and UPR DEGs is found in glucose metabolism, in which genes involved in gluconeogenesis (*Gpt2* and *Pck2*) and UPR (*Atf3*, *Atf4*, *Chop/Ddit3*) are enriched.

**Table 1.**
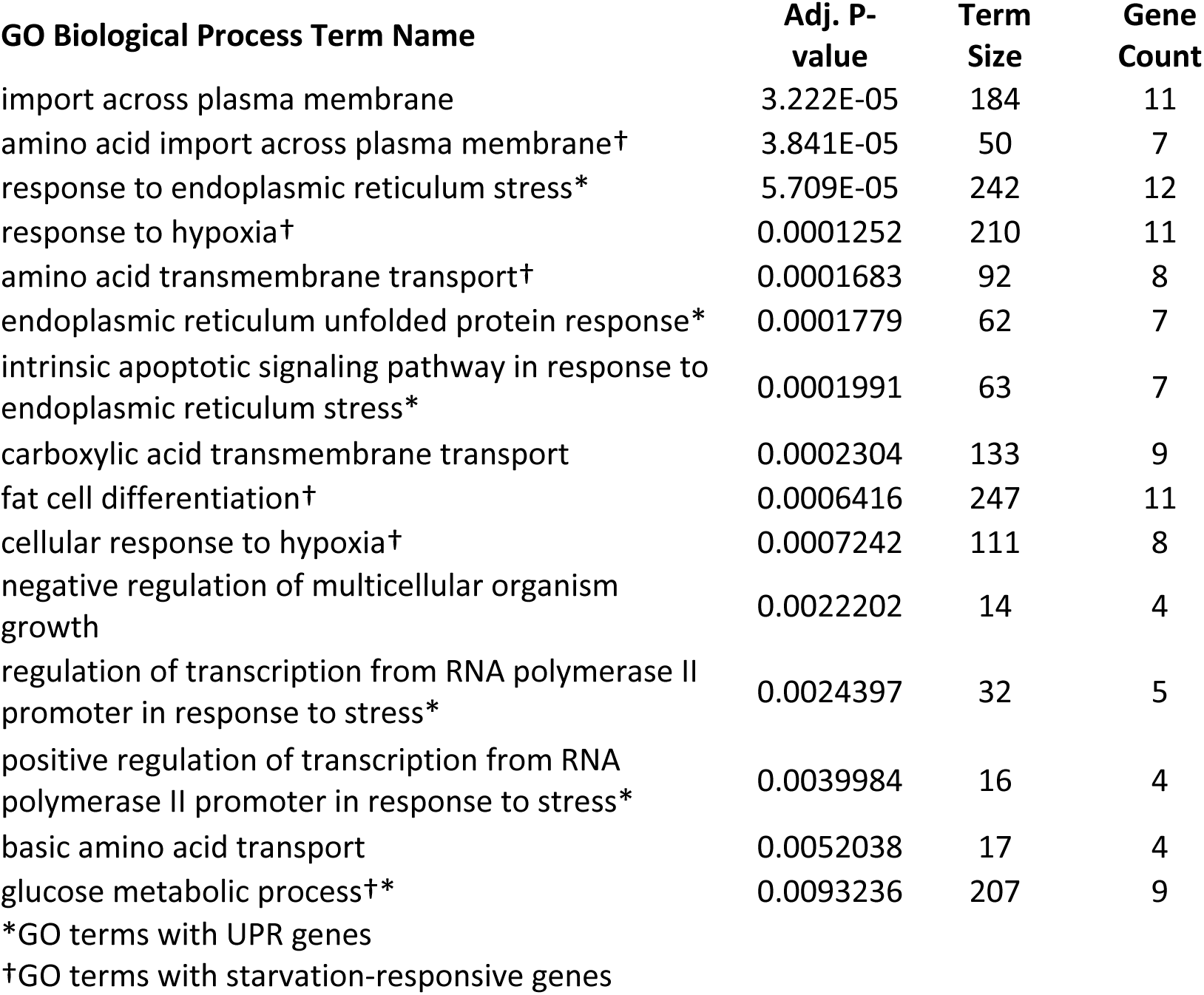
Enriched GO BP terms for upregulated DEGs in *FicD*^F/F^ starved/ *FicD*^F/F^ unstressed.

Next, we compared the DEGs from *FicD*^F/F^ MEFs at 2-hour and 4-hour glucose refeeding with starved and 2-hour refed, respectively. Upon 2-hour refeeding, the *FicD*^F/F^ cells showed the largest number of DEGs, highlighting a large transcriptional response upon the reintroduction of glucose. Many of the UPR genes upregulated during glucose starvation are downregulated at 2- hours of glucose refeeding, suggesting a recovery from the starvation induced UPR (**Fig. 4B**). At 4-hours of glucose refeeding, the remainder of the UPR genes (*Atf3* and *Ero1a*), as well as an additional UPR gene (*Thbs1*) that was upregulated at 2-hour refeeding, are lowered towards unstressed levels (**Fig. 4C**).

Upregulation of UPR genes during glucose starvation, followed by downregulation of UPR DEGs during recovery highlights the homeostatic nature of the response to metabolic stress. This observation is further confirmed by the inverse correlation of DEGs from 2-hour glucose-refed MEFs (compared to starved cells) to DEGs from 4-hour glucose-refed MEFs (compared to 2-hour glucose-refed cells) (**Fig. 4D**). One exception in *FicD*^F/F^ MEFs is ATF3; it remains slightly upregulated at 2-hours of glucose refeeding in wild-type cells. Taken together, our RNA sequencing analysis is consistent with *FicD*^F/F^ MEFs undergoing metabolic stress and PERK responsive UPR transcription during glucose starvation and undergoing recovery during refeeding.

### Transcriptional response to glucose starvation is muted in *FicD^-/-^* MEFs

In contrast to the enhancement of the UPR genes during glucose starvation in *FicD*^F/F^ MEFs (**Fig. 4A**), differential gene expression is almost absent in glucose starved *FicD*^-/-^ MEFs when compared to unstressed *FicD*^-/-^ cells (**Fig. 4E**). DEGs are restricted to 11 genes being upregulated and 6 genes being downregulated, with a notable absence of altered UPR gene expression. Although the trend towards lower expression levels for the UPR genes analyzed by RT-qPCR *FicD*^-/-^ MEFs is still observed in the RNA seq data, these genes do not significantly stand out among total transcripts.

To ensure this loss of response was not an artifact of our RNA seq analysis with EdgeR, we compared the EdgeR-defined DEGs to those defined by additional methods (DESeq2, NOISeq, and limma). All four methods exhibited an overlapping gene expression profile for both *FicD*^F/F^ and *FicD*^-/-^ MEFs (**Fig. S3A** and **B**). The intersection of DEGs defined by all methods represents only 3 genes, compared to 186 intersecting DEGs in *FicD*^F/F^ MEFs. The top two downregulated genes in *FicD*^-/-^ MEFs were *Txnip* and *Arrdc4* (**Fig. 4E**), which play roles in suppressing glucose uptake into cells (*37*). These two genes are downregulated to a similar extent in the *FicD*^F/F^ MEFs upon glucose starvation. Taken together, the data suggests a loss of differential expression in response to glucose starvation in *FicD*^-/-^ MEFs without directly influencing the maintenance of glucose homeostasis.

### Transcriptional response to glucose refeeding is robust in *FicD^-/-^* MEFs

Though the *FicD*^-/-^ MEFs exhibit a muted transcriptional response during glucose starvation, significant changes in the transcriptional profile are observed in the *FicD*^-/-^ MEFs upon glucose refeeding. Like the transcription response observed for *FicD*^F/F^ MEFs (**Fig. 4B)**, the *FicD*^-/-^ MEFs exhibit most DEGs after 2-hour glucose refeeding (**Fig. 4F**). In fact, many of the genes that are upregulated in response to the reintroduction of glucose overlap in the two genotypes (352 overlapping genes represent 58% of upregulated *FicD*^F/F^ MEFs and 68% of upregulated *FicD*^-/-^ MEFs genes, **Fig. S3C**). Given this overlap, we observed a similar inverse correlation of 2-hour refeeding with 4-hour refeeding for *FicD*^-/-^ MEFs genes (**Fig. 4H**). Additionally, downregulated genes in the *FicD*^-/-^ MEFs upon glucose refeeding (*Crebrf*, *Chac1*, and *Parp16*) overlap with downregulated UPR genes in the *FicD*^F/F^ MEFs. However, the *Chac1* downregulation in *FicD*^-/-^ MEFs is dampened (40% of starved levels in *FicD*^-/-^ MEFs vs. 4% of starved levels in *FicD*^F/F^ MEFs) due to its selective upregulation in starved *FicD*^F/F^ MEFs. Chac1 encodes the glutathione-specific gamma-glutamylcyclotransferase 1 responsible for glutathione depletion and the pro-apoptotic effects of the Atf4-Atf3-Ddit3/chop cascade, which is suppressed in starved *FicD*^-/-^ MEFs.

### Transcriptional UPR response to glucose starved and refeeding is aberrant in *FicD^-/-^* MEFs

Because UPR genes were noticeably diminished in glucose starved *FicD*^-/-^ MEFs, we used PCA and heatmap clustering to examine expression patterns of UPR-specific genes in *FicD*^F/F^ and *FicD*^-/-^ MEFs during glucose starvation and refeeding. PCA of the UPR-specific genes highlights altered expression levels for *FicD*^F/F^ MEFs during starvation and refeeding (**Fig. 4J**). However, for *FicD*^-/-^ MEFs, gene expression response to glucose starvation is extremely muted, as illustrated by a comparison of the shifts in each genotype depicted by the dotted arrows (**Fig. 4J**).

When examining heat maps of UPR-specific genes, it becomes evident that *FicD*^-/-^ MEFs exhibit either a delayed or dampened UPR response to glucose starvation and refeeding compared to *FicD*^F/F^ MEFs. The UPR genes form three notable clusters (**Fig. 4K**). The first cluster (**Fig 4K**, *1) includes genes regulated by the PERK arm of the UPR (gene names colored red). This cluster exhibits notable elevation of transcript levels specific to the glucose starved *FicD*^F/F^ MEFs including genes that are slightly elevated in *FicD*^-/-^ starved cells. A second cluster **(Fig 4K**, *2) includes UPR associated transcripts that are consistently elevated in *FicD*^-/-^ MEFs but not *FicD*^F/F^ MEFs, regardless of the glucose treatment. The third cluster (**Fig 4K**, *3) of genes is generally elevated in *FicD*^F/F^ MEFs 2-hour glucose refed cells, with a subset being elevated also in starved *FicD*^F/F^ MEFs. Only three of the genes in this cluster also exhibit elevated levels in 2-hour refed *FicD*^-/-^ MEFs with respect to their unstressed counterparts (**Fig. 4K**). In all three clusters of UPR-specific genes and in both *FicD*^F/F^ and *FicD*^-/-^ MEFs, the 4-hour glucose refed state most closely resembled the unstressed conditions, suggesting that MEFs recover from starvation induced UPR by 4-hours of glucose refeeding.

### FicD loss causes fundamental changes in the transcriptome

We hypothesized that the muted transcriptomic response to glucose starvation in *FicD*^-/-^ MEFs might stem from variations in the baseline unstressed transcriptional profile of these cells. To examine the baseline for *FicD*^-/-^ MEFs, we compared unstressed *FicD*^-/-^ MEFs with unstressed *FicD*^F/F^ MEFs (**Fig 3C**, *FicD*^-/-^/*FicD*^F/F^ genotype response). As previously suggested by the transcriptome PCA (**Fig 3B**), a substantial count of DEGs emerged when comparing the unstressed cells of these two genotypes (**Fig 5A)**, with 852 genes upregulated and 656 genes downregulated in *FicD*^-/-^ MEFs.

**Figure 5.**
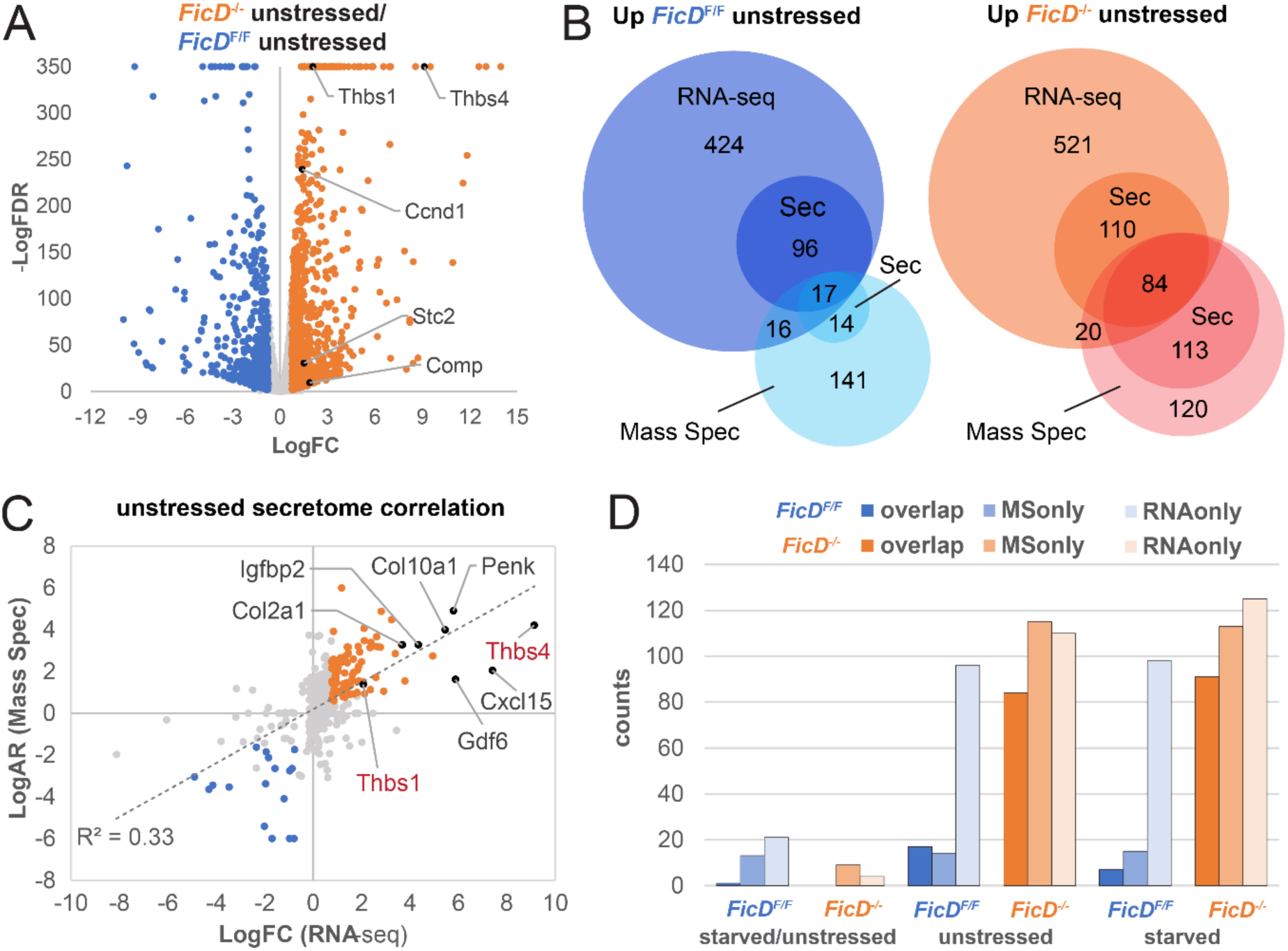
Genotype comparisons reveal upregulation of secretome in *FicD*^-/-^ MEFs. **A)** Volcano plot highlights RNA-seq genotype comparisons of *FicD*^-/-^ unstressed with respect to *FicD*^F/F^ unstressed MEFs. Nodes represent DEGs upregulated in *FicD*^-/-^ (orange), upregulated *FicD*^F/F^ (blue), or other genes (gray). UPR genes are labeled (black). **B)** Venn diagrams compare all and secreted subset of RNA-sec defined DEGs (RNA) with all and secreted subset of significantly different Mass Spec (MS) identified proteins for unstressed *FicD*^F/F^ / unstressed *FicD*^-/-^ MEFs (blue/cyan shades) and unstressed *FicD*^-/-^ MEFs / unstressed *FicD*^F/F^ MEFs (orange/red shades). **C)** Correlation of Log2 fold change (LogFC) for secreted transcriptome (RNA-seq) with Log2 abundance ratio for secreted proteome (MS) for all overlapping genes/proteins in *FicD*^-/-^ unstressed / *FicD*^F/F^ unstressed MEFs. Nodes represent *FicD*^F/F^ upregulated by both RNA and MS (blue), *FicD*^-/-^ upregulated by both RNA and MS (orange), and others (gray). UPR and other top upregulated genes in *FicD*^-/-^ are labeled. **D)** Bar graph depicts secreted gene or protein counts defined by both RNA and MS (dark shades), MS only (medium shades), and RNA only (light shades) for the indicated comparisons.

Among the 852 genes upregulated in *FicD*^-/-^ MEFs, 5 are related to UPR. Only one of these UPR genes, *Stc2*, is also upregulated in the glucose starved *FicD*^F/F^ MEFs (compared to unstressed *FicD*^F/F^ MEFs). *Stc2* encodes a secreted peptide hormone, stanniocalcin-2, whose expression is upregulated by oxidative stress, hypoxia, and the pharmacological stress inducer TG, supporting the prosurvival function of the UPR (*38–40*). Only in the *FicD*^F/F^ starved MEFs is the upregulation of *Stc2* accompanied by the PERK arm of the UPR, suggesting that its baseline upregulation in unstressed *FicD*^-/-^ MEFS is a result of an alternate response, potentially related to hypoxic stress(*41*). Numerous additional hypoxia response genes are differentially regulated between the unstressed *FicD*^F/F^ and *FicD*^-/-^ RNA-seq datasets (102 genes total). A heatmap of their expression levels clusters the genes according to genotype, with roughly one third of the set upregulated specifically in the *FicD*^F/F^ MEFs, one third upregulated specifically in the *FicD*^-/-^ MEFS, and one third upregulated in both genotypes (**Fig. S3D**).

Three of the UPR genes upregulated by genotype comparison in the unstressed *FicD*^-/-^ MEFs (*Thbs4*, *Thbs1*, and *Comp*) encode ECM glycoproteins with thrombospondin type-3 repeats that bind calcium (*42*). Consistent with the roles of these glycoproteins in the ECM, the top enhanced molecular function terms for the DEGs from the *FicD*^-/-^ MEFs that were upregulated with respect to the *FicD*^F/F^ MEFs (**Table 2**) included “extracellular matrix structural constituent”, “glycosaminoglycan binding”, “collagen binding”, and other terms describing secreted protein functions. Taken together, our comparison of the unstressed transcriptional profiles of *FicD*^F/F^ and *FicD*^-/-^ MEFs indicate substantial differences in the gene expression patterns of *FicD*^-/-^ MEFs and an enrichment in secreted proteins in the absence of FicD.

**Table 2.**
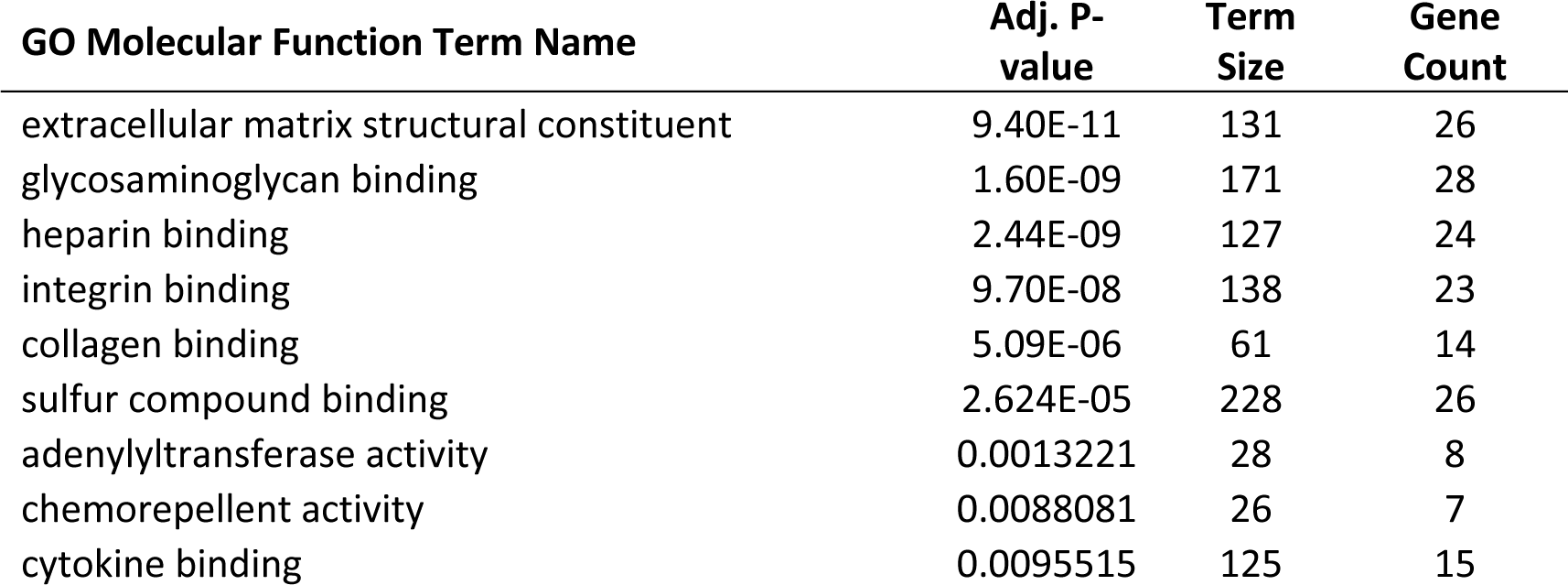
Enriched GO MF terms for upregulated DEGs in *FicD*^-/-^ unstressed/*FicD*^F/F^ unstressed.

### Increased protein secretion in *FicD*^-/-^ MEFs

Given the enrichment of transcripts for secreted proteins among upregulated DEGs in the unstressed *FicD*^-/-^ MEFs (compared to unstressed *FicD*^F/F^ MEFs), we reasoned the absence of FicD and thereby BiP AMPylation may increase the levels of proteins reaching the extracellular space. Previous reports have also suggested that deletion of *FicD* impacts the secretion of cytokines and immunoglobulins in B cells (*43*). To test this idea, we opted to analyze proteins secreted from both genotypes under unstressed and glucose-starved conditions using tandem mass spectrometry (MS/MS).

First, we defined the overlap between the differentially expressed transcripts identified by RNA-seq and the secreted proteome identified by MS/MS by comparing the genotypes grown under unstressed conditions. Among the DEGs that were elevated in *FicD*^F/F^ compared to *FicD*^-/-^ MEFs, approximately one-fifth of the transcripts encoded secreted proteins. Of these, only 17 (3%) were identified as elevated by MS/MS (**Fig. 5B, left panel)**. A relatively larger proportion of the transcripts elevated in *FicD*^-/-^ MEFs compared to *FicD*^F/F^ MEFs encode secreted proteins (26%), and they exhibit a larger overlap (84 proteins, 12%) with the secreted proteins identified by MS/MS (**Fig. 5B, right panel**). Though many of the differentially regulated transcripts observed by RNA seq analysis do not overlap with the altered proteins seen by MS/MS analysis (**Fig. 5B**), a positive correlation (R^2^ = 0.33 or R = 0.57) exists between transcripts and their overlapping MS/MS identified proteins (**Fig. 5C**). Two of the upregulated UPR transcripts from *FicD*^-/-^ MEFs, Thbs1 and Thbs4, were also detected as secreted proteins by MS/MS.

Next, we compared the secretomes of starved MEFs with to those of the unstressed MEFs. *FicD*^F/F^ starved cells exhibit slightly elevated levels of secreted proteins and transcripts when compared to unstressed conditions, with little overlap. These levels are both lower in the *FicD*^-/-^ MEFs, which have no overlap between identified secreted proteins and their transcripts (**Fig 5D, starved/unstressed**). These lower levels reflect the muted transcription response of *FicD*^-/-^ MEFs in response to starvation (**Fig. 4A**). Despite this muted comparative response, both unstressed and starved *FicD*^-/-^ MEFs have elevated secretomes when compared to *FicD*^F/F^ MEFs under the same conditions (**Fig 5D, orange vs. blue**).

## Discussion

BiP is a key player in maintaining protein homeostasis within the ER, and its reversible AMPylation by FicD enzyme has been implicated in modulating its chaperone activity in response to ER stress. The findings presented in this study shed light on the relationship between BiP AMPylation and the UPR response to physiological stress induced by glucose starvation and refeeding. Unlike the ER stress-inducing drugs TG and DTT, glucose starvation led to an induction of PERK responsive UPR transcription that was accompanied by increased BiP AMPylation. Both the UPR response to glucose starvation and BiP AMPylation decreased in *FicD*^F/F^ MEFs upon reintroduction of glucose (**Fig. 1C-E**). This surprising inverse response of BiP AMPylation to chemical and physiological treatments suggests that different stresses induce distinctive outcomes in modulating the amount of active BiP and the UPR response. Chemical inducers of ER stress are pleotropic, do not necessarily reflect UPR resulting from unfolded proteins, and many times are irreversible and therefore not easily resolved (*34, 44*). By contrast, deprivation of glucose in MEFs more closely simulates a reversible physiological ER stress. The inverse upregulation of AMPylation under starved conditions emulated the PTM status of BiP in CHX treated cells with short exposure to DTT stress (**Fig 1B**). For starvation, it is tempting to speculate that the increase in BiP AMPylation reflects an ER stress-initiated decrease in protein translation (akin to CHX treatment) to reduce the energetic cost of ER folding and overcome the lower availability of sugar substrates for protein glycosylation.

The lack of BiP AMPylation in *FicD*^-/-^ MEFs led to a muted transcription response to glucose starvation and refeeding as measured by qPCR for a few known UPR genes (**Fig. 2C**). To gain a more comprehensive understanding of the impact of FicD on UPR-specific and overall gene expression under metabolic stress, we performed RNA-seq analysis of *FicD*^F/F^ and *FicD*^-/-^ MEFs grown in different glucose treatment conditions. The glucose starved *FicD*^F/F^ cells upregulate transcripts for amino acid import, hypoxia response, and glucose metabolism (**Table 1**). Initiation of these biological processes suggests the MEFs tolerate glucose starvation by switching their energy source to amino acids, as has been observed in cells exposed to hypoxia (*45*). The reported hypoxia induced glucose tolerance depended on 5ʹ-AMP-activated protein kinase (AMPK), a well-known stress response activated by glucose starvation. AMPK restores cellular energy balance during stress by sensing intracellular levels of AMP, ADP and ATP (*45, 46*). The kinase is activated by mitochondrial stressors that increase the AMP or ADP to ATP ratio in cells. FicD activity could conceivably alter these nucleotide levels during the AMPylation and deAMPylation cycles of BiP, providing a mechanism for ER stressors to influence AMPK and the energetic status of the cell. Glucose starvation of the *FicD*^F/F^ MEFs also increased expression of genes from the PERK arm of the UPR (**Fig. 4A,K**), which is consistent with reported relationships between low glucose levels and induction of the PERK-CHOP pathway (*47, 48*). The UPR induction was muted in the starved *FicD*^-/-^ MEFs, which exhibited limited transcriptional responses to starvation when compared to unstressed cells. This differential transcription response suggests that FicD is essential for mediating a basal response to glucose starvation in MEFs.

The baseline gene expression profile of unstressed *FicD*^-/-^ MEFs differed substantially from that of *FicD*^F/F^ MEFs (**Fig. 5A**), and perhaps this leads to the lack of response to glucose starvation in these cells. Numerous genes encoding secreted ECM and glycosaminoglycan binding proteins were transcriptionally upregulated in the AMPylation-deficient cells (**Table 2**). Notably, some of these genes are also associated with the UPR. *Thbs4* transcript was upregulated 560-fold in *FicD*^-/-^ MEFs from a very low baseline in *FicD*^F/F^ MEFs, while *Thbs1* and *Comp* transcripts were upregulated 4.2-fold and 3.7-fold, respectively. Thbs4 and Thbs1 function as adhesive glycoproteins in cell-to-cell and cell-to-matrix interactions but also interact with ATF6 in the ER during stress to activate the UPR (*49*). The Comp protein typically functions in the ECM, but mutations of the *Comp* gene can cause intracellular retention of the protein and activation of the apoptotic arm of the UPR, resulting in skeletal dysplasia (*50, 51*). Overall, the lack of FicD appears to deregulate the expression of genes and in unstressed cells an elevation of expression is observed for multiple DEGs (Fig. 5 and Table 2).

A single UPR gene, Stanniocalcin-2 (Stc2), was upregulated in both the glucose-starved *FicD*^F/F^ MEFs and the unstressed *FicD*^-/-^ MEFs. The Stc2 gene encodes a glycosylated peptide hormone that functions as a pro-survival component of the UPR and negatively modulates store-operated Ca^2+^ uptake (*39, 40*). In response to traditional UPR-inducing drugs tunicamycin and thapsigargin, Stc2 expression is upregulated through the PERK/ATF4 mediated pathway, together with other noted UPR markers (e.g., CHOP/Ddit3_1 and Herpud1) (*40*). The glucose starved *FicD*^F/F^ MEFs activated similar PERK pathway genes as the tunicamycin/thapsigargin treated cells. However, the unstressed *FicD*^-/-^ MEFs activated *Stc2* in the absence of an inducer. This *Stc2* upregulation in the unstressed *FicD*^-/-^ MEFs suggests the inability to AMPylate BiP causes an alternate or premature ER stress. *Stc2* is also upregulated in response to hypoxia and oxidative stress, and many of the genes that respond to hypoxia are differentially regulated in the RNA-seq dataset (Fig. S2D). Among numerous hypoxia response genes whose differential regulation is limited to *FicD*^F/F^ MEFs, *Eif4ebp1* is upregulated in both starved and 2-hour glucose refed states. *Eif4ebp1* is a target of the ATF4 transcription factor and represses translation initiation in response to oxidative stress (*52, 53*). The dual activation of eIF2α through the UPR and eIF4e through oxidative stress in *FicD*^F/F^ MEFs would reduce translation to levels where BiP is not required for chaperoning and is therefore AMPylated by FicD (*54*).

Expanding upon our observation of the baseline upregulation of transcripts for extracellular proteins in *FicD*^-/-^ MEFs compared to *FicD*^F/F^ MEFs, we conducted an analysis of protein secretion using tandem mass spectrometry. While the secretome transcript and protein levels correlated, their overlap between the two detection methods remained low in the MEF genotype comparisons (Fig 5). This low overlap could result from many factors. The abundance of secreted proteins represents a balance of protein synthesis, turnover, and travel through the secretory pathway, that would differ in *FicD*^F/F^ and *FicD*^-/-^ MEFs. The detection of proteins is dependent on the fidelity of detection for a specific peptide and therefore some proteins are overrepresented while others are underrepresented. Despite these limitations, our MS/MS analysis revealed that the abundance of secreted proteins were elevated in *FicD*^-/-^ MEFs when compared to *FicD*^F/F^ MEFs. The comparison of mRNA levels and MS/MS identification of secreted proteins suggest that *FicD*^-/-^ MEFs secreted a significantly greater quantity of proteins than *FicD*^F/F^ MEFs in both unstressed and starved conditions.

Because inhibition of translation is a primary consequence of UPR signaling, the elevated secretion in *FicD*^-/-^ cells may reflect an inability to constrain protein synthesis. The increased secretome levels in AMPylation-deficient MEFs accompany an increase in protein synthesis observed under all glucose treatment conditions. Thus, protein secretion appears to become dysregulated in *FicD*^-/-^ MEFs where the rheostat for BiP has been deleted. Even under conditions of starvation stress, which usually represses protein translation (*55*), the hypersecretion response remained unaffected in MEFs lacking FicD. The impaired translation regulation in *FicD*^-/-^ MEFs was also observed in the transcription profile of pre-ribosome genes that function in ribosome biogenesis (Fig. S3E). Genes that were upregulated upon glucose refeeding in *FicD*^F/F^ MEFs, were enriched in the cellular component GO term ‘pre-ribosome’. Most of these genes follow a similar expression pattern trend. They are highest in *FicD*^F/F^ MEFs upon 2-hours of feeding and persist at lower levels through 4-hours of refeeding. At the same time, their levels are decreased in *FicD*^-/-^ unstressed and starved conditions and elevated slightly after 2-hours of refeeding, but not after 4-hours. These results provide strong evidence that, in the ER, FicD plays a crucial role in regulating cellular stress responses, transcriptome changes, and the composition of secreted proteins.

Our observations, along with previous studies, propose that the elevated UPR and delayed recovery of the UPR in FicD knockout cells and tissues may be attributed to the excessive chaperoning effect exerted by the heightened active pool of BiP in the absence of FicD. However, the absence of a discernible ERAD gene response in *FicD^-/-^* MEFs suggests that the elevated UPR and delayed recovery in the absence of FicD may originate primarily from the abundant differentially expressed genes in the baseline *FicD^-/-^*MEFs rather than being solely attributed to excessive chaperone activity of BiP. This indicates a complex interplay of factors influencing the UPR dynamics in the absence of FicD, where both altered gene expression and the chaperoning effect of BiP may contribute to the observed cellular responses. Overall, these observations support the proposal that a response by the ER is not isolated to the environment of the ER but integrates signals from the cell, similar to previous proposals for an Integrated Stress Response (IRS) (*56*).

In summary, our study provides compelling evidence for the role of FicD in modulating cellular responses to both pharmacological and physiological ER stress. The absence of FicD leads to alterations in gene expression patterns, disruption of UPR dynamics, abnormal secretion of proteins, and dysregulation of translation; shedding light on the intricate interplay among FicD, ER stress, and cellular homeostasis. These findings emphasize the complexity of the UPR and its sensitivity to diverse stressors. To date, the only validated substrate for FicD mediated AMPylation is BiP. FicD is proposed to act as molecular rheostat for BiP activity. The ability of FicD to control amounts of active BiP by AMPylation and deAMPylation adds additional fidelity to how cells response to UPR. As observed previously, metazoans, such as flies and mice, require this rheostat in tissues composed of differentiated cells essential for an animal’s lifetime, such as the eye and pancreas, respectively (*28, 30*). The presence of this rheostat is necessary to safeguard these tissues from lifelong stress cycles. Further research is needed to elucidate the precise mechanisms through which FicD’s impact extends beyond ER stress and to investigate the functional significance of the observed gene expression changes in the context of ER stress and the more far-reaching IRS (*56*). Our study provides valuable insights into the molecular pathways governing cellular responses to physiological stress and highlights potential genes and pathways for therapeutic interventions in diseases associated with dysregulated ER stress.

## Materials and Methods

### Reagents and general remarks

All the reagents were purchased from Thermo Fisher Scientific and Sigma unless otherwise stated and were of appropriate grade. Antibodies were purchased from Abcam, Cell Signaling, and Thermo Fisher Scientific except the monoclonal α-AMP antibody which is a generous gift of Aymelt Itzen (University Medical Center Hamburg Eppendorf, Germany). Unless otherwise stated, all the qPCR experiments were repeated at least three times and the western blot experiments repeated at least twice, by using distinct samples.

### Isolation and immortalization of MEFs

Isolation: *FicD*^F/F^ and *FicD*^-/-^ mouse embryos (E13.5) were cleaned from placenta and membrane in PBS in a 10 cm petri dish under dissection microscope. Embryos were broken by sucking and exhausting 5 times with 2.5 mL syringe with 18G needle. Each broken embryo placed into 10 cm dish in 10 mL complete medium: High-Glucose DMEM medium (Sigma, D5796) supplemented with 10% FBS (Sigma, F2442), 1 mM sodium pyruvate (Sigma), 1x penicillin/streptomycin-L-glutamine (Sigma). Cells were grown in 5% CO_2_ at 37 °C for 2 days, large bones and unbroken pieces were removed by sedimentation, and split 1/3 ratio. Splitting and sedimentation were repeated every 2-3 days total 3 times and cells were collected by centrifugation and frozen with 10% DMSO/complete medium by slow cooling (20 °C 1h to 80 °C) in cryotubes. Immortalization: Thawed MEFs were split into 6 well plates by ¼ and 1/6 dilutions and grown overnight in 5% CO_2_ at 37 °C. Approximately 25% confluent wells were transfected with 2 µg plasmid vector expressing SV40 antigen using Fugene Transfection Reagent (Promega) as described by manufacturer. MEFs were incubated overnight in 5% CO_2_ at 37 °C and medium was exchanged. Two days after transfection, transfected MEFs were split into 10cm dish. MEFs were split to ultimately 1/100,000 fold dilution by observing the confluence over the course of 3-4 passages to eliminate non-transformed cells. Immortalized MEFs were frozen by slow cooling and stored in liquid nitrogen.

### Cell culture

MEFs were grown to 80–90% confluency in standard 10cm cell culture dish (VWR) and cultured in complete medium: High-Glucose DMEM medium (Sigma, D5796) supplemented with 10% FBS (Sigma, F2442), 1 mM sodium pyruvate (Sigma), 1x penicillin/streptomycin-L-glutamine (Sigma). MEFs were incubated at 5% CO2 at 37 °C. Pharmacological treatment of MEFs were carried out in complete medium supplemented with 0.1-10 µM thapsigargin in DMSO or 5mM DTT in PBS or 100 µg/mL CHX. Glucose starvation was carried out in glucose-free medium: Glucose-free DMEM (Thermo Fisher, 11-966-025) supplemented with 10% FBS (Sigma, F2442), 1x penicillin/streptomycin-L-glutamine (Sigma).

### Cell Harvesting and Lysis

MEFs were grown in 6-well plates in 5% CO_2_ at 37 °C to 80% confluence. Treated MEFs were either harvested by trypsinization or transferred to ice and harvested by scraping and transferred to 1.5 mL reaction tubes on ice. Medium were aspirated after centrifugation at 1000xg for 5 min at 4 °C and MEFs were washed with ice cold PBS. After aspirating the PBS, MEFs were frozen in liquid nitrogen and stored at 80 °C or directly lysed with lysis buffer (50 mM HEPES pH 8.0, 150 mM NaCl, 0.5% NP40, 0.5% Triron-X100, 5% glycerol, 1x Roche Complete EDTA free inhibitor cocktail, 1x Roche PhosSTOP) for 1h on ice by agitation. Lysed MEFs were centrifuged at full speed for 10 min on a bench top Eppendorf 5424 R centrifuge at 4 °C and supernatants were transferred to fresh reaction tubes. Lysate concentrations were determined by Pierce BCA Protein Assay (Thermo Scientific). For mRNA isolation, frozen MEFs were lysed directly with QIAshredder columns (Qiagen).

### Western-blotting

Normalized lysate samples were resolved with 12% SDS-PAGE gels using Mini-PROTEAN® Tetra Cell (Bio-Rad) and blotted to poly(vinylidene difluoride) membrane for 10 min at 1.5 Amps using a Power Blotter (Invitrogen). The membranes were washed with TBS-T buffer and blocked with 5% BSA or 1x Roti-Block (Carl Roth) for 1 h at room temperature. Primary antibody was directly added and incubated overnight at 4 °C (α-AMP (gift from Aymelt Itzen), α-GRP78 (Abcam, ab21685), α-eIF2α-phospho (Abcam, ab32157), α-eIF2α (Cell Signaling, 9722)). After washing, secondary horseradish peroxidase (HRP) conjugated antibodies (goat α-mouse (Abcam, ab205719), donkey α-rabbit (Amersham, NA934)) with 1:5,000-10,000 dilution respectively were added and incubated for 1 h at room temperature. The membranes were washed and visualized using ChemiDoc Imaging System (Bio-Rad) after incubation with Pierce ECL Western Blotting Substrate (Thermo Scientific) for 2–3 min.

### RNA isolation, RT-qPCR, and RNA-seq

Total RNA from MEFs were isolated by RNeasy Plus Mini Kit (Qiagen) as described by the manufacturer after lysing the MEFs by QIAshredder columns (Qiagen). RNA concentrations were measured by nanodrop (Thermo Scientific). Normalized RNA samples were digested with DNase I (Thermo Scientific) in 20 µL reaction (2 µg total RNA) for 30 min at 37 °C. DNase reaction was stopped by adding 2 µL of EDTA (50mM) at 65 °C for 10 min. DNase treated RNA was reverse transcribed by qScript cDNA synthesis kit (Quanta Bio) in a 40 µL reaction as descried by the manufacturer. Quantitative PCR was performed with 10 ng cDNA by using PowerTrack SYBR Green Master Mix (Thermo Scientific) in either 96 well plates or 384 well plates using either CFX 96 (Bio-Rad) or CFX Opus 384 (Bio-Rad) respectively. U36B4 (NM_007475) was used as the reference mRNA. Results were analyzed by CFX Maestro Software (Bio-Rad) and plotted by Prism 9 (Graphpad). Primers used for the genes analyzed in this study (U36B4 forward primer: 5’ cgtcctcgttggagtgaca 3’, reverse primer: 5’ cggtgcgtcagggattg 3’; ATF3 forward primer: 5’ tggagatgtcagtcaccaagtct 3’, reverse primer: 5’ gcagcagcaattttatttctttct 3’; ATF4 forward primer: 5’ actctaatccctccatgtgtaaagg 3’, reverse primer: 5’ caggtaggactctgggctcat 3’; CHOP forward primer: 5’ ccagaaggaagtgcatcttca 3’, reverse primer: 5’ actgcacgtggaccaggtt 3’; sXBP1 forward primer: 5’ctgagtccgcagcaggt 3’, reverse primer: 5’ tgtcagagtccatgggaaga 3’; BiP forward primer: 5’ caaggattgaaattgagtccttctt 3’, reverse primer: 5’ ggtccatgttcagctcttcaaa 3’; FicD forward primer: 5’ gtagacgcactgaatgagttcg 3’, reverse primer: 5’ tggtgtataagtagtcagcctgg 3’. For RNA-seq experiment, isolated RNAs are first passed the quality control then sequenced by Novogene Corporation Inc. and raw data is mapped with CLC Genomics Workbench software (version 9.5, CLC Bio, Aarhus, Denmark).

### Analysis of RNA-seq

Fastq reads corresponding to each of four treatments (unstressed, glucose starved, 2-hour refed, and 4-hour refed) applied to *FicD*^F/F^ and *FicD*^-/-^ MEFs (all conditions in triplicate) were mapped to the mouse reference genome, and statistical analysis was performed using CLC Genomics Workbench software (version 9.5, CLC Bio, Aarhus, Denmark). Total counts and CPM for each mouse gene were generated for all conditions (8 conditions in triplicate, 24 samples). Principal Component Analysis (PCA) was calculated for all samples with ClustVis (*57*) using Log2 CPM counts with unit variance scaling and row centering. Principal components were calculated using Singular Value Decomposition (SVD) with imputation.

Total counts mapped for each sample were used to calculate differential gene expression with four methods: EdgeR, DESeq2, limma, and NOISeq using the Integrative Differential Expression Analysis for Multiple EXperiments (IDEAMEX) server with chosen parameters (LogFC=0.75, FDR=0.05, and CPM=1) without batch effects (*58*). Differentially expressed genes (DEGs) were identified for various pairwise conditions (**Fig. 2C**). Differential expression results were integrated to observe the consistency of DEGs identified by each method **(Fig. S3A** and **B**). Ultimately, EdgeR (*59*) DEGs were chosen for subsequent GO term enrichment.

For GO term enrichment analysis, Ensemble IDs for upregulated EdgeR DEGs were submitted to the G:profiler server (*60*) limiting the statistical domain scope to annotated genes, using the G:SCS threshold to calculate adjusted P-values with a cutoff below 0.01, limiting pathway size to between 10 and 250 terms, and excluding electronic GO annotations. We removed redundant terms with identical lists of genes, keeping the top term ranked by the lowest P-value. We report the enriched GO biological process terms for upregulated DEGs from metabolic stress comparison of *Fic*^FL/FL^ starved/*FicD*^F/F^ unstressed MEFs and for enriched GO molecular function terms for upregulated DEGs from genotype comparison of *FicD*^-/-^ with *FicD*^F/F^ MEFs.

PCA and heat maps were generated for mouse DEGs (identified an any of the RNA-seq comparisons from Fig3C, as well as DEGs from any of the conditions compared to the respective unstressed state) with GO terms related to unfolded protein response (UPR) as defined in UniProt (*61*). Genes were clustered in the heatmap by hierarchical clustering (Euclidean distance with Ward method) using clustvis with the same parameters as used for PCA (*57*).

### Preparation of Secretomes

After overnight incubation in serum-containing media with or without glucose, the culture media of MEFs were replaced with serum-free media with or without glucose. Following an additional incubation period of 2-4-hours, the supernatants from the MEFs were collected into 1.5 mL tubes placed on ice. To remove cells and debris, the supernatants were centrifuged for 15 minutes at 3200 x g at 4 °C. Subsequently, the supernatants were filtered through a 0.22μm filter into fresh microcentrifuge tubes. To the filtered supernatants, a final concentration of 150 μg/ml sodium deoxycholate was added and incubated for 15 minutes on ice. Furthermore, a final concentration of 8% (v/v) trichloroacetic acid was added to each sample, followed by an overnight incubation at 4 °C. The precipitated proteins were resuspended and transferred to fresh microcentrifuge tubes that had been pre-rinsed with methanol to remove collagen contamination. These suspensions were then subjected to centrifugation for 1 hour at 27,000 x g at 4 °C. Afterward, the supernatants were discarded, and the pellets were washes twice with 1.5 ml of pre-cooled 100% acetone, with each wash involving centrifugation for 40 minutes at 27,000 x g at 4 °C. Following the second wash, the pellets were air-dried for 10 minutes and then resuspended in 1 ml of 10 mM Tris-HCl at pH 8.0. The resuspended samples were transferred to fresh microcentrifuge tubes in preparation for tryptic digestion.

### Tandem Mass Spectrometry

Secreted protein samples were reduced with 10mM DTT for 1 hr at 56°C and alkylated with 50mM iodoacetamide for 45 min at room temperature in the dark. Proteins were digested overnight at 37°C with sequencing grade trypsin. Resulting peptides were then de-salted via solid phase extraction (SPE) prior to analysis. LC-MS/MS experiments were performed on a Thermo Scientific EASY-nLC 1200 liquid chromatography system coupled to a Thermo Scientific Orbitrap Fusion Lumos mass spectrometer. To generate MS/MS spectra, MS1 spectra were first acquired in the Orbitrap mass analyzer (resolution 120,000). Peptide precursor ions were isolated and fragmented using high-energy collision-induced dissociation (HCD). The resulting MS/MS fragmentation spectra were acquired in the ion trap. Label-free quantitative searches were performed using Proteome Discoverer 2.2 software (Thermo Scientific). Samples were searched against all reviewed entries in the Mouse UniProt protein database. Searches included the following modifications: carbamidomethylation of cysteine residues (+57.021 Da), oxidation of methionine (+15.995 Da), and acetylation of peptide N-termini (+42.011 Da). Precursor and product ion mass tolerances were set to 10 ppm and 0.6 Da, respectively. Peptide spectral matches were adjusted to a 1% false discovery rate (FDR) and proteins were filtered to a 5% FDR. All samples were run in biological triplicate.

### Analysis of Secretomes

Proteins were mapped from the following treatment and genotype comparisons: starved *FicD*^F/F^/ unstressed *FicD*^F/F^, starved *FicD*^-/-^ / unstressed *FicD*^-/-^, unstressed *FicD*^-/-^ / unstressed *FicD*^F/F^, and starved *FicD*^-/-^ / starved *FicD*^F/F^. Upregulated proteins were defined as having an abundance ratio fold change = >1.5 and p-value < 0.05, as well as proteins with missing values that were identified in at least 2 reps of the first condition (i.e. starved *FicD*^F/F^ from starved *FicD*^F/F^ / unstressed *FicD*^F/F^) but 0 reps of the second (i.e. unstressed *FicD*^F/F^ from starved *FicD*^F/F^ / unstressed *FicD*^F/F^). Proteins with low combined FDR confidence (<0.05) were excluded. The same cutoffs were used to identify downregulated proteins (abundance ratio = <1.5 and p-value < 0.05, as well as proteins with missing values that were identified in at least 2 reps the second condition but 0 reps of the first condition). Secreted proteins were identified from UniProt annotations for GO cellular component terms that include “extracellular” or for subcellular location terms that include “secreted”.

## Dataset S1

### RNA sequencing data and functional enrichment

Sheet 1, dataset condition summary. Sheet 2, RNA sequencing differential expression data. Sheet 3, Average Log2CPM of UPR, Hypoxia, and Ribosome Biogenesis genes. Sheet 4, differential expression for secreted proteins. Sheet 5, functional enrichment for *FicD*^F/F^ starved/*FicD*^F/F^ unstressed MEFs. Sheet 6, functional enrichment for *FicD*^F/F^ 2 hr/*FicD*^F/F^ starved MEFs. Sheet 7, functional enrichment for *FicD*^-/-^ unstressed/*FicD*^F/F^ unstressed MEFs.

## Dataset S2

### Mass Spec Data

Sheet 1, *FicD*^-/-^ unstressed/*FicD*^F/F^ unstressed MEFs. Sheet 2, *FicD*^-/-^ starved/*FicD*^F/F^ starved MEFs.

## Acknowledgments

We extend our gratitude to Aymelt Itzen at the University Medical Center Hamburg Eppendorf, Germany, for generously providing the monoclonal α-AMP antibodies. We appreciate the assistance provided by various UT Southwestern Labs, including the Peter Douglas Lab in mapping the RNA-seq data, the Jenna Jewell Lab for supplying the plasmid required for SV40 immortalization, Michael Buszczak for critical reviews and the Jun Wu Lab for sharing the MEF isolation protocol. We also acknowledge the invaluable contributions of the Orth Lab members for their insightful discussions, editing, and support. KO is a WW Caruth, Jr. Biomedical Scholar with an Earl A Forsythe Chair in Biomedical Science. Funding sources include Welch Foundation grant I-1561 (KO), Once Upon a Time…Foundation (KO) and the National Institutes of Health Grant R35 GM134945 (KO). The funders had no role in study design, data collection and interpretation, or the decision to submit the work for publication.

