## Supplementary material for "FicD Sensitizes Cellular Response to Glucose Fluctuations in Mouse Embryonic Fibroblasts": Supp Figs

Fig. S1.

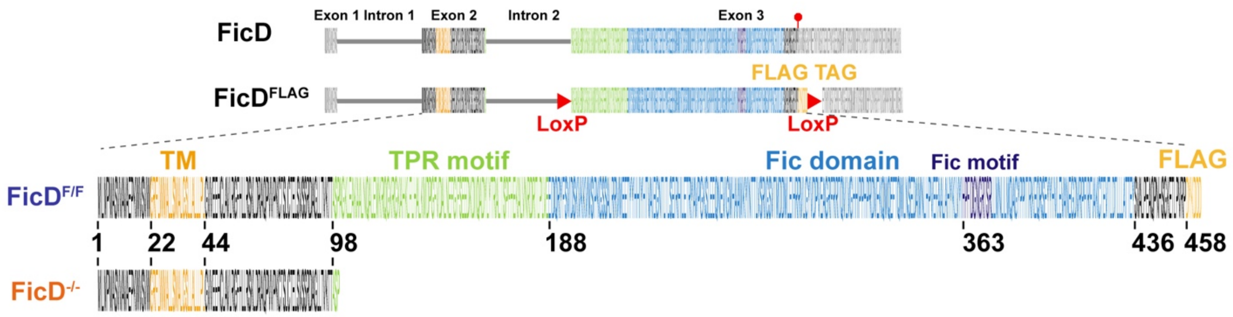


**Figure S1. FicD knockout mouse embryonic fibroblasts (MEFs).** FicD-FLAG and FicD knockout MEFs: *FicD*$^{F/F}$ and *FicD*$^{-/-}$. Insertion of FLAG tag to wild type MEFs and deletion of Fic domain and TPR motif from LoxP flanked *FicD*$^{F/F}$ MEFs is shown.

Fig. S2.

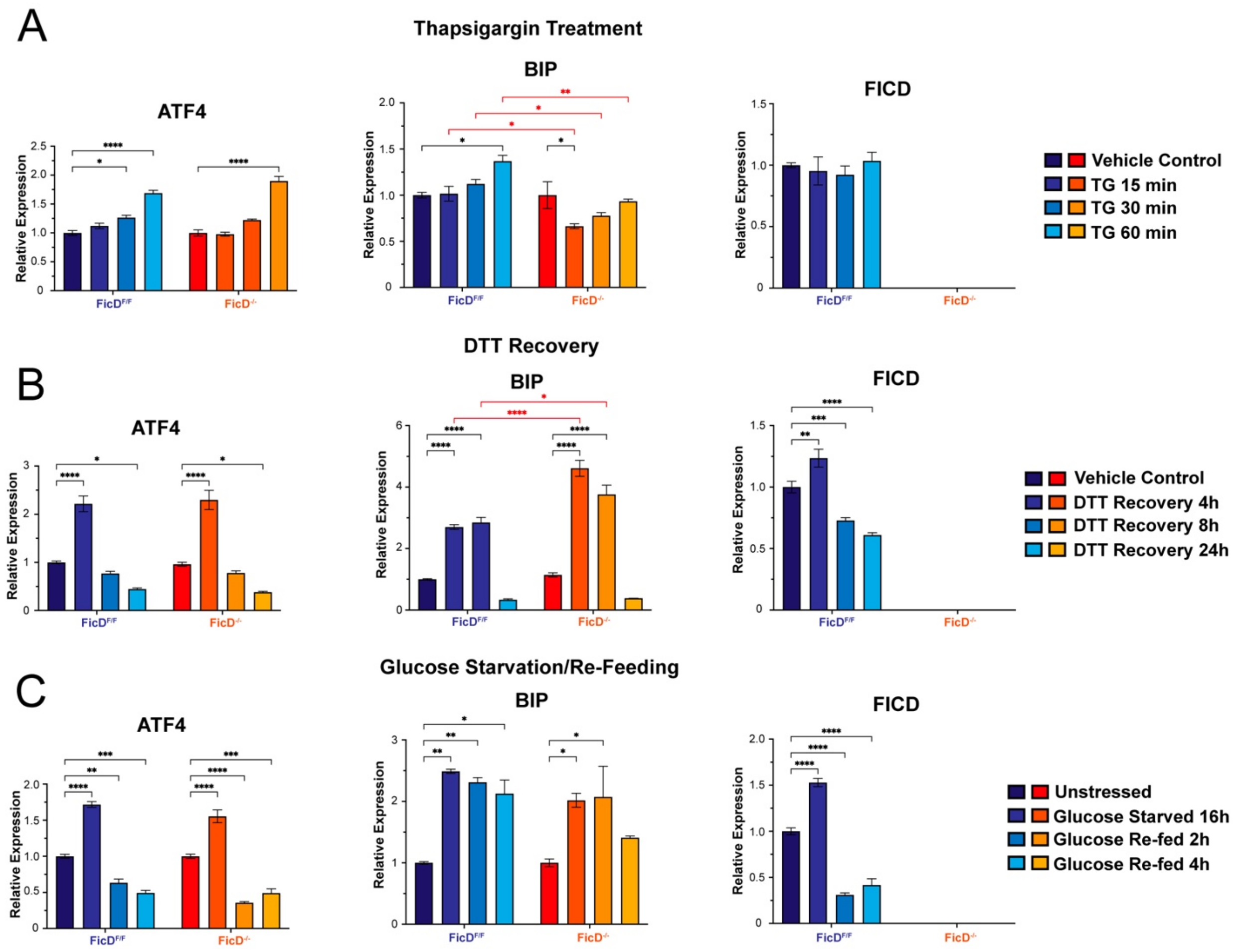


**Figure S2. Thapsigargin (TG), DTT, and Glucose Starvation induced ER stress of *FicD*$^{F/F}$ and *FicD*$^{-/-}$ MEFs.** RT-qPCR showing relative expression levels of UPR marker genes ATF4, BiP, and FicD upon **A)** treatment of MEFs with Thapsigargin (1μM), or **B)** recovery of MEFs from 1h DTT (5mM) exposure, or **C)** glucose starvation (for 16h) and re-feeding of MEFs for indicated time points. Error bars represent the standard deviation of 3 biologically independent repeat of the experiment with 4 technical replicates. Genotypes are demonstrated by shades of blue for *FicD*$^{F/F}$ and shades of orange for *FicD*$^{-/-}$ MEFs. Two-way ANOVA with Tukey multiple comparison test is applied to determine the significance. P values: 0.1234 (ns), 0.0332 (*), 0.0021 (**), <0.0001 (****). Non-significant comparisons are not shown for clarity.

**Fig. S#3**

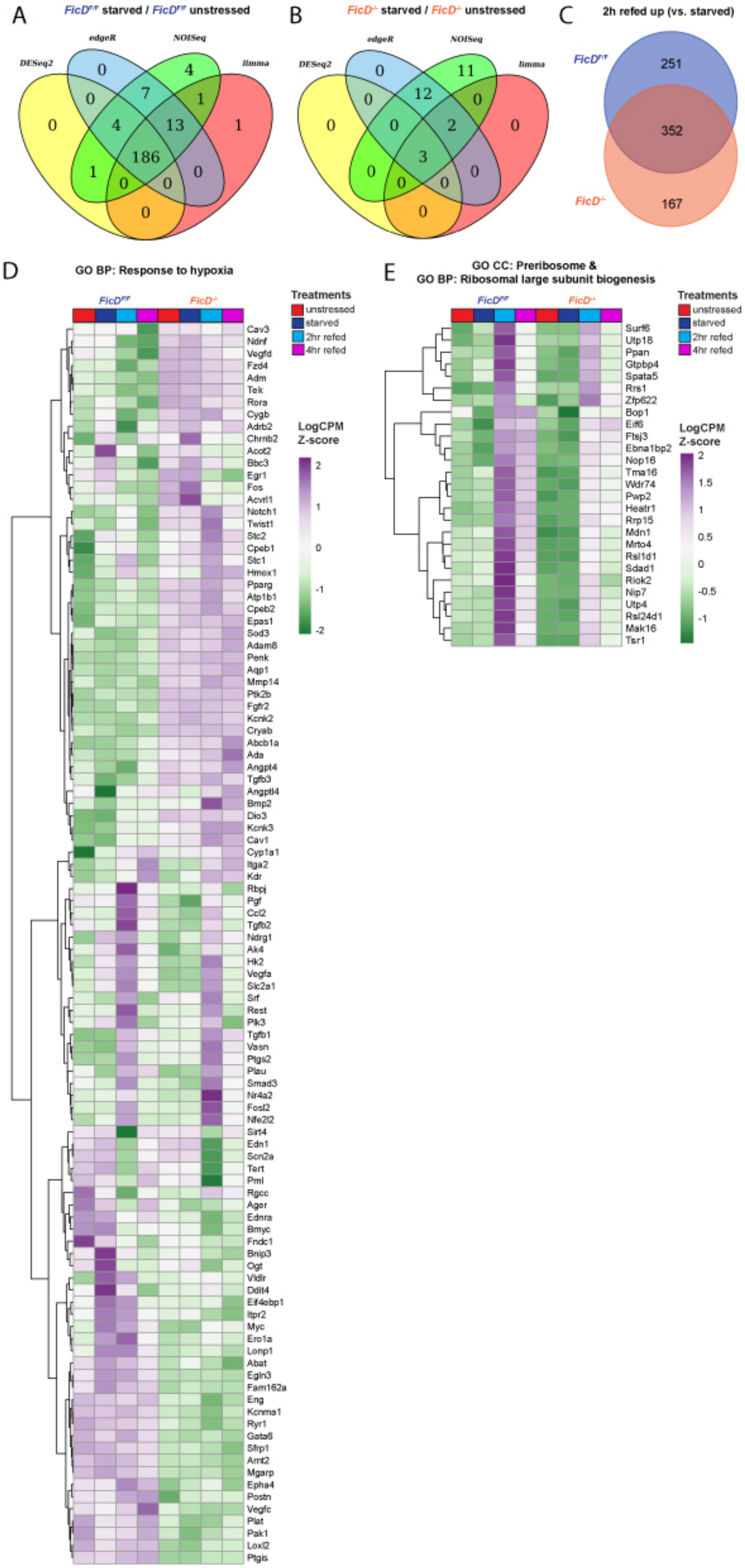
A
FicD^f/f starved / FicD^f/f unstressed
B
FicD-/- starved / FicD-/- unstressed
C
2h refed up (vs. starved)
251
352
167
D
GO BP: Response to hypoxia
E
GO CC: Preribosome &
GO BP: Ribosomal large subunit biogenesis
Treatments
unstressed
starved
2hr refed
4hr refed
LogCPM
Z-score

**Figure S3. RNA-seq comparisons of DEGs defined by different methods** for A) *FicD*$^{F/F}$ starved / *FicD*$^{F/F}$ unstressed MEFs and B) *FicD*$^{-/-}$ starved / *FicD*$^{-/-}$ unstressed MEFs. C) Venn diagram comparing upregulated genes in both genotypes for 2h refed / starved MEFs. D-E) Heatmap clustering Log2CPM (Z-score) of differentially expressed genes under any condition with D) GO BP term response to hypoxia and E) GO CC term preribosome or GO BP term ribosomal large subunit biogenesis.

**Dataset S1 -Supplemental_Dataset_S1 (**separate file**)**

**Dataset S2 -Supplemental_Dataset_S2 (**separate file**)**
